# SARS-CoV-2 infection triggers pro-atherogenic inflammatory responses in human coronary vessels

**DOI:** 10.1101/2023.08.14.553245

**Authors:** Natalia Eberhardt, Maria Gabriela Noval, Ravneet Kaur, Swathy Sajja, Letizia Amadori, Dayasagar Das, Burak Cilhoroz, O’Jay Stewart, Dawn M. Fernandez, Roza Shamailova, Andrea Vasquez Guillen, Sonia Jangra, Michael Schotsaert, Michael Gildea, Jonathan D. Newman, Peter Faries, Thomas Maldonado, Caron Rockman, Amy Rapkiewicz, Kenneth A. Stapleford, Navneet Narula, Kathryn J. Moore, Chiara Giannarelli

## Abstract

COVID-19 patients present higher risk for myocardial infarction (MI), acute coronary syndrome, and stroke for up to 1 year after SARS-CoV-2 infection. While the systemic inflammatory response to SARS-CoV-2 infection likely contributes to this increased cardiovascular risk, whether SARS-CoV-2 directly infects the coronary vasculature and attendant atherosclerotic plaques to locally promote inflammation remains unknown. Here, we report that SARS-CoV-2 viral RNA (vRNA) is detectable and replicates in coronary atherosclerotic lesions taken at autopsy from patients with severe COVID-19. SARS-CoV-2 localizes to plaque macrophages and shows a stronger tropism for arterial lesions compared to corresponding perivascular fat, correlating with the degree of macrophage infiltration. *In vitro* infection of human primary macrophages highlights that SARS-CoV-2 entry is increased in cholesterol-loaded macrophages (foam cells) and is dependent, in part, on neuropilin-1 (NRP-1). Furthermore, although viral replication is abortive, SARS-CoV-2 induces a robust inflammatory response that includes interleukins IL-6 and IL-1β, key cytokines known to trigger ischemic cardiovascular events. SARS-CoV-2 infection of human atherosclerotic vascular explants recapitulates the immune response seen in cultured macrophages, including pro-atherogenic cytokine secretion. Collectively, our data establish that SARS-CoV-2 infects macrophages in coronary atherosclerotic lesions, resulting in plaque inflammation that may promote acute CV complications and long-term risk for CV events.

## Main

Coronavirus disease 2019 (COVID-19), caused by severe acute respiratory syndrome coronavirus 2 (SARS-CoV-2), is uniquely marked by extraordinary tissue tropism and an array of clinical presentations from asymptomatic infection to acute respiratory distress, multiorgan failure, and death^1^. Ischemic cardiovascular events such as acute myocardial infarction (AMI) and stroke, due to the underlying disruption of a chronically inflamed atherosclerotic plaque^2^, are established clinical complications of COVID-19^1,3^. AMI and stroke can be triggered by several other acute respiratory viral infections, including influenza virus^4^. However, patients with COVID-19 are > 7-fold more likely to have a stroke than patients with influenza^5^, and their risk for both AMI and stroke remains high for up to 1 year after infection^6^. The extreme inflammatory response that occurs in severe cases of COVID-19, also known as a cytokine storm^7^, is likely a contributor to the increased risk for AMI and stroke. Yet the possibility that SARS-CoV-2 directly affects the coronary vasculature, as documented for other distant organs (e.g., kidney, gut, brain, adipose tissue, and myocardium)^8^, remains largely unexplored. A main aggravator of tissue damage in the lungs is a potent inflammasome activation in macrophages in response to SARS-CoV-2 virus^9^. A similar response in macrophages that infiltrate arterial vessels could boost plaque inflammation and risk for AMI and stroke in COVID-19 patients. Here we show that infiltrating macrophages are infected by SARS-CoV-2 in coronary autopsy specimens from COVID-19 patients. Lipid-laden macrophages (foam cells), a hallmark of atherosclerosis at all stages of the disease^10^, are more susceptible to SARS-CoV-2 infection than other macrophages and this is dependent on the receptor neuropilin-1 (NRP-1). SARS-CoV-2 induces a strong pro-atherogenic inflammatory response in both macrophages and foam cells, which is recapitulated in *ex-vivo* SARS-CoV-2 infection of human vascular explants. This response gives us insight into what may be a contributing factor to the ischemic cardiovascular complications observed in COVID-19 patients.

### Spatial artificial intelligence identifies SARS-CoV-2 viral RNA in macrophages within coronary arteries from deceased COVID-19 patients

We analyzed coronary autopsy specimens (*n* = 27) from 8 patients with RT-PCR confirmed diagnosis of COVID-19 between May 2020 and May 2021. Demographics and clinical characteristics, including past medical history, cardiovascular risk factors and other relevant clinical information, were obtained from the patients’ electronic medical records and autopsy reports (IRB i21-01587) (**Fig. 1A; Supplementary Tables 1,2**). Mean age was 69.6 (median 71; 59-84) and 75% of patients were male (6/8). Patients presented with coronary artery disease (8/8), 3 or more cardiovascular risk factors such as hypertension (8/8), overweight or obesity (7/8), hyperlipidemia (7/8), type 2 diabetes (6/8), and chronic kidney disease (4/8), and some had a history of either myocardial infarction (1/8) or ischemic stroke (1/8) (**Fig. 1A; Supplementary Table 1**). Acute myocardial ischemia was diagnosed during hospitalization in 3 patients, 1 presented stroke, and 4 were found to have coronary stenosis (> 50%) at autopsy. The patients, with the exception of 1 pronounced dead before admission, were hospitalized for an average of 17.6 days (range 8–31) (**Supplementary Table 2**). One patient (Pt. 1), was hospitalized 3 times (for a total of 31 days) following the first diagnosis. Pt.1 developed acute myocardial infarction and died during the third hospitalization approximately 140 days after testing negative for COVID-19 (**Fig. 1A; Supplementary Table 2**). The clinical presentation, in hospital course of the disease, COVID-19 treatments, as well as macroscopic and microscopic autopsy pathology, obtained from hospital medical records and autopsy reports and are summarized in **Supplementary Table 2**.

**Figure 1.**
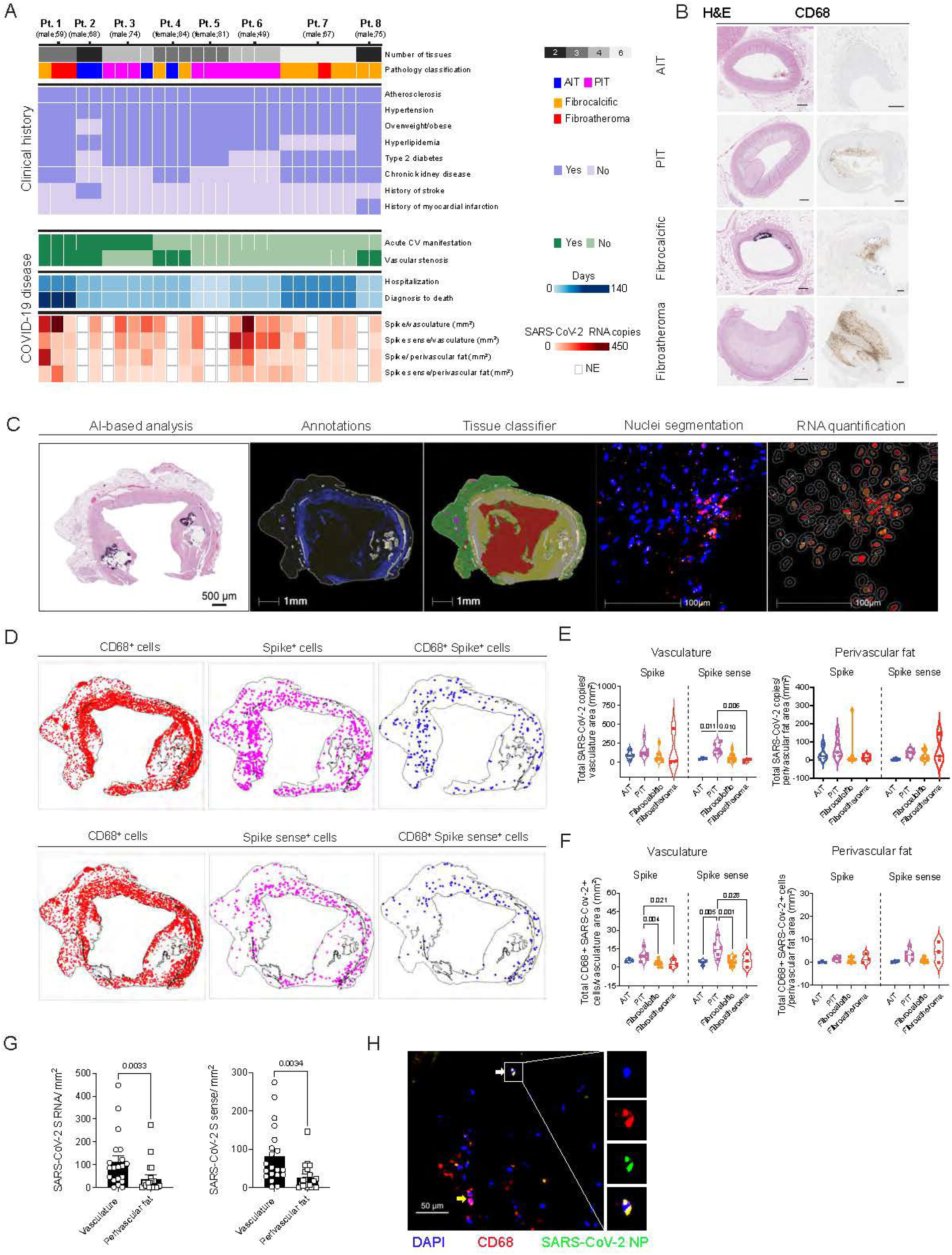
SARS-CoV-2 viral RNA in human coronary arteries from deceased individuals with COVID-19 is identified using artificial intelligence-based spatial analysis. A) Categorical heatmap of coronary autopsy specimens (*n* = 27) from deceased individuals with COVID-19 (*n* = 8) displays their pathology classification into adaptive intimal thickening (AIT), pathological intimal thickening (PIT), fibrocalcific plaques, and fibroatheromas. Clinical history for atherosclerosis, hypertension, overweight/obese status, hyperlipidemia, type 2 diabetes, chronic kidney disease, history of stroke, and myocardial infarction for each patient are also shown. Summary of acute cardiovascular (CV) manifestations during COVID-19 disease progression, coronary stenosis (*no*: <50% or *yes*: >50%), hospitalization duration, and time to death (days) following a first PCR confirmed COVID-19 diagnosis are also depicted. Spike and spike sense RNA copy number normalized to vasculature and perivascular fat area (mm^2^) are depicted for each sample. NE: not evaluated. B) Representative images of coronary human samples stained with hematoxylin and eosin (*left*), and CD68 chromogenic immunohistochemistry staining (*right*) for each pathological classification. Scale = 500 μm. C) Representative images of *in situ* RNA-FISH AI-based analysis. Following semi-automatic annotations, AI-based neural network was used to classify the arterial wall (vasculature, *yellow*) and perivascular fat (*green*). Background and artifacts (*red*) were removed from the analysis. Next, nuclei (DAPI+) segmentation classifier analysis and RNA quantification were performed using HALO AI, HALO, and spatial analysis workflow. D) Representative images of spatial analysis of coronary samples showing the location of the cells that are positive for CD68 (*red dots*), SARS-CoV-2 spike (*magenta dots, top*) or spike sense (*magenta dots, bottom*) cells, and CD68 SARS-CoV-2 RNA double positive cells (*blue*). E) Violin plots showing total SARS-CoV-2 RNA copies of spike and spike sense probes normalized by either vascular wall or perivascular fat area (mm^2^) in AIT, PIT, fibrocalcific, and fibroatheroma coronaries. F) Violin plots showing total CD68+ SARS-CoV-2+ spike or spike sense cells in the vasculature or perivascular fat regions normalized by vasculature or perivascular fat area (mm^2^) in the different pathological classification. ANOVA one-way statistical analysis with post-hoc Tukey’s test for multiple comparisons was performed to evaluate differences between groups. G) SARS-CoV-2 spike (S) and spike sense (S sense) quantification in vasculature and perivascular fat normalized by tissue area (mm^2^). Paired t-test was performed to evaluate differences between two groups. H) Representative images of multicolor immunofluorescence of SARS-CoV-2 nucleoprotein (NP) (*green*) and CD68 (*red*) in human coronary tissues from patient with COVID-19. White arrow indicates CD68+ SARS-CoV-2+ cell and yellow arrow depicts CD68+ cell, DAPI (blue) identify the nuclei. Scale = 50 um

Sections of coronary arteries from all autopsies were stained with hematoxylin and eosin (H&E) and classified by a clinical cardiovascular pathologist (N.N.) as adaptive intimal thickening (AIT; *n* = 4), pathological intimal thickening (PIT; *n* = 10), fibrocalcific plaque (*n* = 10), and fibroatheroma (*n* = 3) (**Fig. 1A,B; Extended Data Fig. 1A,B; Supplementary Table 3**). Detailed pathological features, including presence of lipid pool, necrotic core, and adventitial inflammation, were noted (**Extended Data Fig. 1B**). Macrophage infiltration was identified using immunohistochemical staining for CD68, revealing a larger positive stained area corresponding to the necrotic cores preferentially in fibroatheromas (**Fig. 1B, Extended Data Fig. 1C, Supplementary Table 3**).

To identify SARS-CoV-2 viral RNA (vRNA) in the post-mortem coronary vasculature from patients with COVID-19, we performed RNA fluorescence in-situ hybridization (RNA-FISH) analysis for the viral RNA encoding the spike (S) protein. To establish whether SARS-CoV-2 infected the human coronary vasculature, we also probed the antisense strand of the S gene, which is only produced during viral replication. A CD68 probe was used to identify macrophages infiltrating the coronary vessels in the same sections and to establish the cellular localization of SARS-CoV-2 vRNA. As SARS-CoV-2 infects fat depots and accumulates vRNA in adipose tissue to trigger a strong proinflammatory response^11,12^, we used a neural network artificial intelligence (AI) approach to classify the coronary arterial wall from perivascular fat in each sample while we employed nuclei segmentation to quantify the RNAscope probes in cells infiltrating the two tissues (**Fig. 1C-D**). In the coronary arterial wall, vRNA encoding S protein and the antisense strand of the S gene were detected at different levels in all the sections from all patients, indicating the presence of vRNA and replicative activity of the virus **(Fig. 1E**, vasculature). SARS-CoV-2 S gene copy number was similar across AI, PIT, fibrocalcific, and fibroatheroma coronary lesions. However, PIT coronaries showed a significantly higher copy number of the antisense strand of the S gene (identified by the S sense probe), indicating higher viral replication in the vascular wall of these lesions (**Fig. 1E**). In particular, CD68+ cells expressing both the SARS-CoV-2 S and the antisense strand of S (S sense probe) were significantly higher in the vasculature of PIT coronaries *vs* other pathologies **(Fig. 1F)**. In perivascular fat, vRNA encoding S protein was detected in 20 of the 21 sections and the S sense in 19 of the 21 sections (**Fig. 1A**). Overall, each patient presented at least one section positive for S and S sense vRNA (**Fig. 1A,E,F**). Notably, the amount of S and antisense strand vRNA was significantly lower in perivascular fat than in the corresponding arterial wall across all samples (**Fig. 1G**). The accumulation of viral protein material in the coronaries was confirmed by immunofluorescence (**Fig. 1H**). PIT arterial lesions, which appeared more susceptible to SARS-CoV-2 infection based on the presence of the sense and antisense strands of the S gene, contained significantly more cells than other types of lesions, and 4.8-fold more cells than corresponding perivascular tissue (2691.8 ± 288.7 vs 697.6 ± 159.3 cells/ mm^2^; *P* < 0.0001) (**Extended Data Fig. 1D**). The significantly higher number of CD68 RNA+ cells in the coronary vasculature in both PIT and fibroatheromas (**Extended Data Fig. 1E**), corresponded to a higher number of CD68 RNA^+^ cells in perivascular fat of PIT lesions than fibrocalcific lesions and similar to fibroatheromas (**Extended Data Fig. 1E**). This suggests SARS-CoV-2 directly infects perivascular macrophages to increase coronary susceptibility to SARS-CoV-2 infection. This possibility was further supported by the significant association between SARS-CoV-2 vRNA copies with CD68 copy number in both arterial wall and perivascular fat (**Extended Data Fig. 1F-G**). Notably, more SARS-CoV-2 vRNA encoding S protein accumulated in the vascular and perivascular fat tissue (*P* = 0.06) and the coronary wall (*P* = 0.08) from COVID-19 patients with acute cardiovascular manifestations than in patient without CV (**Extended Data Fig. 1H**).

### SARS-CoV-2 infection of foam cells and macrophages

The accumulation of cholesterol-laden macrophages (foam cells) is a hallmark of atherosclerosis at all stages of the disease from early PIT to late fibroatheroma lesions^10,13^. To investigate SARS-CoV-2 infection of both macrophages and foam cells, we differentiated human monocytes derived from human peripheral blood mononuclear cells (PBMCs) into macrophages, and treated them with oxidized low-density lipoprotein (LDL) complexed with Dil dye (Dil-Ox-LDL) to differentiate them into foam cells. To experimentally confirm our observation that SARS-CoV-2 can infect human plaque macrophages, macrophages and foam cells were infected either with the icSARS-CoV-2 mNeonGreen (mNG) reporter virus, a modified virus that uses mNG fluorescence as a surrogate readout for viral replication^14^, or SARS-CoV-2 USA WA1/2020 isolate. mNG expression confirmed the ability of SARS-CoV-2 to replicate in both cell types, although replication was significantly higher in foam cells (**Fig. 2A; Extended Data Fig. 2A**). We confirmed that foam cells are more susceptible to the virus as significantly more nucleoprotein (NP) accumulated in foam cells compared to macrophages following infection with SARS-CoV-2 USA WA1/2020 isolate (**Fig. 2B**; **Extended Data Fig. 2B**). In fact, although the number of SARS-CoV-2 NP^+^ foam cells fell between 24- and 48-hpi, they still remained proportionally higher than the number of SARS-CoV-2 NP+ macrophages (**Fig. 2B**). Foam cells also accumulated more SARS-CoV-2 S vRNA than macrophages (**Fig. 2C**). The SARS-CoV-2 vRNA genome was detectable in both macrophages and foam cells as early as 2 hpi, remained high up to 24 hpi, but was reduced at 48 hpi in both cell types (**Fig. 2D; Extended Data Fig. 2D**), although the reduction was greater in macrophages than in foam cells (**Fig. 2E**). This decay in vRNA levels suggests that although they are both susceptible, neither macrophages nor foam cells can sustain a productive viral infection. To test this explicitly, we performed a plaque assay using modified Vero E6 cells expressing the transmembrane protease serine 2 and human angiotensin-converting enzyme 2 (Vero E6-TMPRSS2-T2A-ACE2). The plaque assay confirmed a progressive decline of viral titer in conditioned media from infected macrophages and foam cells over 48 hr (**Fig. 2F, Extended Data Fig. 2C**).

**Figure 2.**
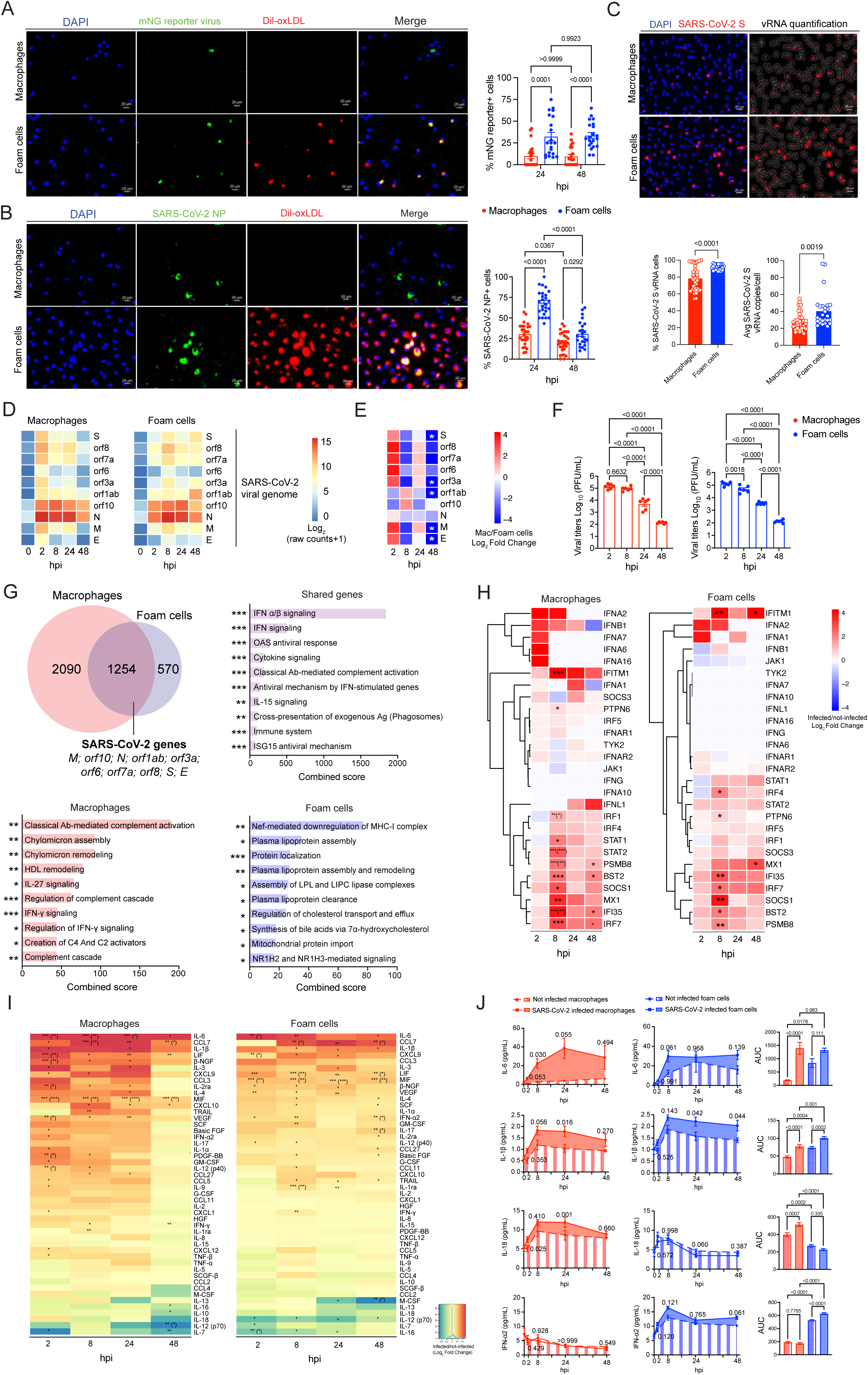
A) Representative fluorescence microscopy images of human primary macrophages and Dil-Ox-LDL laden macrophages (foam cells) cultured with mNG reporter (MOI 0.1) at 48 hours post-infection (hpi). Images depict nuclei with DAPI (*blue*), mNG reporter virus (*green*), Dil-oxLDL treatment (*red*). Scale bar = 20 μm. Bar dot plot quantification of mNG positive macrophages (*red*) or foam cells (*blue*) at 24 and 48 hpi. B) Representative fluorescent images of human primary macrophages and Dil-Ox-LDL laden macrophages (foam cells) infected with SARS-CoV-2 USA WA1/2020 (MOI 0.1) at 48 hpi. Images depict nuclei with DAPI (*blue*), Nucleoprotein (NP; *green*), Dil-Ox-LDL (*red*). Scale bar = 20 μm. Bar dot plot quantification of NP positive macrophages (*red*) or foam cells (*blue*) at 24 and 48 hpi. C) RNA-FISH quantification of frequency and SARS-CoV-2 viral RNA (vRNA) copies using AI-based nuclei segmentation in SARS-CoV-2 infected human primary macrophages and foam cells. SARS-CoV-2 S probe (*red*), DAPI+ nuclei (*blue*). Scale bar = 20 μm. D) Heatmap of SARS-CoV-2 viral reads (Log_2_ scaled raw counts +1) in macrophages and foam cells at baseline (0), 2, 8, 24 and 48 hpi. *S:* spike gene*; N:* nucleocapsid gene*; M:* membrane gene*; E:* envelope gene*; orf:* open reading frames. E) Heatmap of log_2_ fold change of SARS-CoV-2 viral genes in macrophages vs. foam cells at 2, 8, 24 and 48 hpi as determined by bulk RNAseq. Adjusted *P*-values < 0.05 (FDR 1%) were considered significant. F) Viral titer quantification (Log_10_ PFU/mL) by plaque assay in VERO E6-TMPRSS2-T2A-ACE2 cells of culture supernatants from human primary macrophages and foam cells cultured with SARS-CoV-2 USA WA1/2020 (MOI 0.1). ANOVA one-way statistical analysis followed by Tukey’s post-hoc test was performed for multiple comparisons. *P*-values < 0.05 were considered significant. G) Venn diagram of DEGs in infected *vs* non-infected macrophages (*red*; *n* = 2090 genes) and foam cells (*blue*; *n* = 570 genes), and shared genes (*purple*; *n* = 1254 genes). Bar plots show corresponding upregulated signaling pathways (Reactome 2022) in infected macrophages and/or foam cells, ranked by their combined score in *Enrichr*. *\*, P < 0.05; **, P < 0.01; ***, P < 0.001*. H) Heatmaps of Log_2_ Fold change of interferon response genes in SARS-CoV-2 infected macrophages (*left*) and foam cells (*right*) vs. non-infected cells. *P*-values were adjusted using Benjamini-Hochberg (BH) correction (FDR = 10%). Adjusted *P* values < 0.05 were considered significant. Asterisk indicates an adjusted *P* < 0.05 for the comparison of infected vs not infected at each timepoint. Asterisk in parenthesis (*) indicates an adjusted *P value* < 0.05 for the interaction between infection and timepoint terms of the model. *\*, P < 0.05; **, P < 0.01; ***, P < 0.001*. I) Heatmap of 48 cytokines and chemokines secreted from SARS-CoV-2 infected macrophages and foam cells at different times post-infection. Data are shown as Log_2_ fold change vs. mock uninfected cells at each timepoint. *\*, P < 0.05; **, P < 0.01; ***, P < 0.001*. Adjusted *P* values (BH method) in parenthesis. J) Kinetic plots show the area under the curve (AUC) of proatherogenic cytokines secreted by SARS-CoV-2 infected macrophages (*red*) and foam cells (*blue*) versus non-infected cells (*n* = 8) at different times post-infection. One-way ANOVA statistical analysis following Dunnett’s multiple comparisons was performed. Bar plots show the quantification of the AUC for each cytokine. One-way ANOVA statistical analysis following Tukey’s multiple comparisons was performed.

### SARS-CoV-2 drives proatherogenic inflammatory responses in macrophages and foam cells

Based on our observation that SARS-CoV-2 replication was abortive in macrophages and foam cells and existing evidence that an over-reactive inflammatory response to SARS-CoV-2 is orchestrated by macrophages in other tissues^9,12,15^, we investigated the immune response of macrophages and foam cells to SARS-CoV-2. Differential gene expression analysis of RNA-seq data from infected macrophages and foam cells identified shared and unique transcriptional signatures (**Fig. 2G**). As expected, the 1254 shared genes included the SARS-CoV-2 viral genes. Other commonly upregulated genes were those involved with antiviral responses and SARS-CoV-2 infection including interferon (IFN) signaling pathways and antiviral processes by type I and II IFN signaling, OAS antiviral response, negative regulation of viral replication and viral life cycle, as well as complement activation and cytokine signaling. ISG15 antiviral signaling, which dampens IFN signaling and is implicated in the hyperinflammatory response associated with COVID-19 severity^15,16^, was also upregulated in both cell types (**Fig. 2G**; **Extended Data Fig. 2E**).

Infected macrophages expressed a unique transcriptional signature associated with classical complement cascade activation, complement cascade and C4 and C2 activators, IFN-γ signaling and its regulation, and IL-27 signaling, which induces IFN/STAT1-dependent genes^17^, and regulation of cytokine pathways. Genes involved in lipid metabolism were also upregulated, indicating lipid metabolic reprogramming occurs in macrophages in response to the virus (**Fig. 2G; Extended Data Fig. 2E**). The unique transcriptional signature of infected foam cells included 570 genes (**Fig. 2G; Extended Data Fig. 2E**). The top upregulated signaling pathway involved Nef-mediated downregulation of MHC-I, a response induced by viruses to evade immune recognition^18,19^. Infected foam cells also upregulated processes and signaling pathways involved in the regulation of lipid metabolism, that may facilitate viral entry and replication^20^.

We confirmed the upregulation of several genes in the type I IFN response pathway in both SARS-CoV-2-infected macrophages and foam cells, with the strongest response observed at 8 hpi (**Fig. 2H; Extended Data Fig. 2F**). In macrophages, significantly upregulated genes included *IRF1*, a transcriptional activator of IFN alpha and beta, as well as genes induced by IFN alpha, beta, and gamma; *MX1,* encoding the GTP-binding protein Mx1 that has antiviral activity against RNA viruses including SARS-CoV-2 ^21,22^, and *STAT1* and *STAT2. IRF7*, known to induce type I IFN responses, and the viral restriction factor *IFITM1*^23^ were also upregulated in SARS-CoV-2-infected macrophages. SARS-CoV-2-infected foam cells had similar IFN response activation, but a delayed upregulation of *MX1* that only occurred at 48 hpi, consistent with the higher viral RNA and protein accumulation seen in foam cells. Only foam cells upregulated *IRF4,* which inhibits MyD88 signaling and is expressed in M2-like macrophages^24^. Moreover, the expression of *STAT1* and *STAT2* were not significantly increased in foam cells (**Fig. 2H**), suggesting infected macrophages induce a distinct IFN-induced JAK/STAT signaling pathway. Notably, both cell types significantly upregulated several pro-inflammatory and proatherogenic cytokine and chemokine genes (**Extended Data Fig. 2F**), including *CCL7, TNFSF10* (also known as *TRAIL*), *CXCL10, CCL7,* and *CCL3.* Infected macrophages uniquely upregulated *CXCL9*, *CXCL12*, and *CLEC11A,* whereas foam cells upregulated genes included *TNFA*, *CCL5*, and *CCL2*.

To further investigate the inflammatory profile of macrophages and foam cells in response to SARS-CoV-2 infection, we quantified the secretion of cytokines and chemokines for up to 48 hpi in conditioned media. Several pro-inflammatory and pro-atherogenic cytokines (e.g., IL-6, CCL7, IL-1β, β-NGF, IL-3, LIF, MIF, CXCL-9, IFN-α, and IFN-γ) were released following infection of both macrophages and foam cells (**Fig. 2I**). Among these cytokines are ones known to trigger ischemic cardiovascular events, including IL-6, a candidate therapeutic target in ongoing clinical trials^25^, and IL-1β, whose inhibition reduces secondary cardiovascular events in high-risk post-myocardial infarction patients^26^ (**Fig. 2J**). Moreover, the release of macrophage migration inhibitory factor (MIF), a pro-atherogenic and inflammatory cytokine implicated in intima-media thickening, lipid deposition, and plaque instability^27^, was increased in both cell types after infection (**Extended Data Fig. 2G**).

Other cytokines were differentially secreted by macrophages and foam cells in response to SARS-CoV-2 infection, suggesting distinct inflammatory responses exist between the two cell populations. For example, the proatherogenic cytokine IL-18^28^ was significantly released by infected macrophages but not foam cells, whereas IFN-α2, a type I IFN response cytokine which inhibits viral replication^29^, was significantly released by infected foam cells but not macrophages (**Fig. 2J**). Importantly, we detected significant differences in the release of many of these cytokines over time between uninfected macrophage and foam cells, suggesting that differences in the baseline inflammatory status of each cell type influences their response to the virus (**Fig. 2J; Extended Data Fig. 2G**).

### SARS-CoV-2 infection of human atherosclerosis vascular explants enhances a pro-inflammatory plaque microenvironment

To determine whether the proinflammatory response to SARS-CoV-2 we observed in human macrophage and foam cells *in vitro* also occurs in vascular tissue, we infected human atherosclerotic vascular explants with SARS-CoV-2 (USA WA1//2020) and assayed for transcript expression by bulk RNAseq (**Fig. 3A**). SARS-CoV-2 vRNA encoding the structural S, envelope (E), membrane (M), and nucleocapsid (N) proteins, as well as ORFs encoding nonstructural accessory proteins, were detectable in infected plaques as early as 24 hpi and persisted to 72 hpi (**Fig. 3B**). S and N protein expression and virus-like particles were observed in infected atherosclerotic plaques (**Extended Data Fig. 3A,B**). However, viral titer decreased over time with no detectable infectious particles by plaque assay at 72 hpi, suggesting abortive replication in the vascular explants (**Fig. 3C**), similar to what we observed with foam cells and macrophages *in vitro*. Regardless, SARS-CoV-2 infection induced a strong type I IFN transcriptional response in infected plaques reflected by the early upregulation of transcription factors and genes involved in response to viral infections such as *IRF7*, *JAK1*, *IFITM1* at 2hpi; *IFNAR*, *IRF4*, *IRF1, MX1, PTPN6, IFNA1, STAT1* and *STAT2* at 24 hpi; *IRF1*, *IFNA7*, *IFI35* at 48 hpi, and *IFNA16* at 72hpi (**Fig. 3D**). Genes involved in negative regulation of IFN signaling (i.e., *SOCS1* and *SOCS3*) were also upregulated, likely reflecting the activation of regulatory signaling (**Fig. 3D**). Interestingly, SARS-CoV-2 infection of vascular explants triggered the expression of viral receptors and entry factors such as *ACE2, NRP1, FURIN*, *TMPRSS4, TMPRSS11A and CTSB* at 24 hpi, suggesting that the virus facilitates its own entry in host cells (**Fig. 3E**).

**Figure 3.**
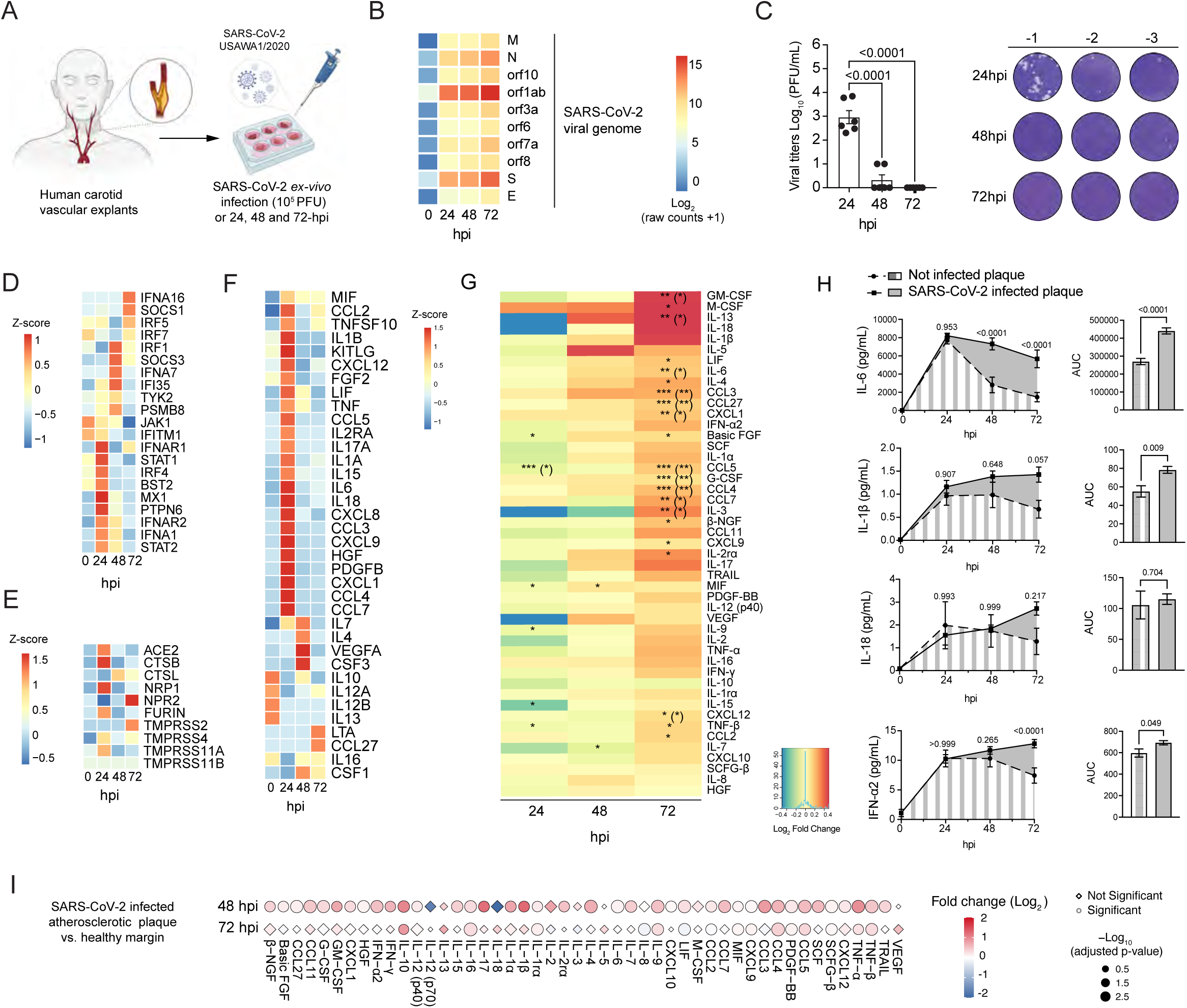
A) Schematics of experimental approach of human carotid vascular explants infection with SARS-CoV-2 USA WA1/2020 (10^5^ PFU/mL). Samples were collected at baseline, 24, 48, and 72 hours post-infection (hpi). B) Heatmap of SARS-CoV-2 viral reads (Log_2_ scaled raw counts +1) in carotid vascular explants at baseline (0), 24, 48, and 72 hpi. *S:* spike gene*; N:* nucleocapsid gene*; M:* membrane gene*; E:* envelope gene*; orf:* open reading frames. C) Infectious viral titer quantification by plaque assay in VERO E6-TMPRSS2-T2A-ACE2 cells of culture supernatants of human carotid plaques infected with SARS-CoV-2 USA WA1/2020. D) Heatmap shows the standardized z-scored expression of interferon response genes in human carotid vascular samples infected with SARS-CoV-2 virus at different times post-infection. E) Heatmap of standardized z-scored expression of selected host viral receptors and entry factors in SARS-CoV-2 infected human carotid vascular samples at baseline, 24, 48, and 72 hpi. F) Heatmap of standardized z-scored gene expression of cytokine and chemokine genes in SARS-CoV-2 infected human carotid vascular explants at different times post-infection. G) Heatmap of cytokines and chemokines secreted from SARS-CoV-2 infected human atherosclerotic plaques at 24, 48, and 72 hpi. Data are shown as log_2_ fold change *vs.* not infected samples. *\*, P < 0.05; **, P < 0.01; ***, P < 0.001*. *P* values in parentheses were adjusted using the Benjamini-Hochberg method. H) Kinetic plots show the area under the curve (AUC) of proatherogenic cytokines and chemokines released in the supernatant of non-infected or SARS-CoV-2 infected carotid plaques at different timepoints (hpi). One-way ANOVA statistical analysis following Dunnett’s multiple comparisons was performed and *P* values are denoted for each timepoint comparison. Bar plots show AUC, paired *t*-test was performed to compare the two groups. I) Plot showing the relative expression of secreted cytokines and chemokines between SARS-CoV-2 infected atherosclerotic plaque tissue vs. SARS-CoV-2 infected vascular margins at 48 and 72 hpi. Relative expression is represented in log_2_ fold change colored scale. Statistical significance is expressed as dot size (-Log_10_ adjusted p-value). Statistically significant values are represented as circles, while not significant changes are represented as diamonds.

SARS-CoV-2 also initiated a transcriptional proinflammatory response that largely recapitulated the one seen in cultured macrophages and foam cells. This included the upregulation of proatherogenic cytokines such as *IL1B, IL6, MIF, ILB, TNF*, *IL7, CCL5*, as well as chemokines like *CCL2, CCL3, CCL4, CCL5, CXCL9, CCL27, CCL7, CCL6, CXCL1, CXCL8, CXCL9,* and *CXCL12*. Anti-inflammatory cytokines, such as *IL10* and *IL13,* were down regulated, further supporting a strong proatherogenic inflammatory response to SARS-CoV-2 infection in human atherosclerotic plaques (**Fig. 3F; Extended Data Fig. 3C**). Analysis of the secretome of infected plaques revealed similar proinflammatory protein changes (**Fig. 3G**). SARS-CoV-2 infected plaques released several proatherogenic cytokines and chemokines including IL-6, IL-1β, and IFN-α2, as well as CCL2, CCL3, CCL4, and CCL7 (**Fig. 3G**). The release of several cytokines and chemokines was significantly higher at 72 hpi. However, only IL-6, IL-1β, IFN-α2, and CCL3 were secreted at significantly higher amounts, calculated as area under the curve (AUC), over time (**Fig. 3H**; **Extended Data Fig. 3D**). A stronger inflammatory response was seen in SARS-CoV-2 infected atherosclerotic plaques versus non-atherosclerotic paired surgical margins, as shown by the significantly higher release of several cytokines (e.g., IFN-γ, IFN-α2, IL-1β, IL-17, TNF-α, TNF-β, CCL3, CCL4, CCL7) from infected plaques, mainly at 48 hpi (**Fig. 3I**). These findings suggest that SARS-CoV-2 infection triggers a hyperactivated immune response mainly within atherosclerotic lesions, a response that could contribute to the increased risk of ischemic cardiovascular events in COVID-19 patients with underlying atherosclerosis.

### NRP1+ foamy macrophages promote atherosclerotic plaque tissue susceptibility to SARS-CoV-2

To elucidate the mechanism of vascular susceptibility to SARS-CoV-2 infection, we evaluated the expression of the main viral entry receptors and co-factors in the aorta, coronary, and tibial arteries using gene expression data publicly available from the Genotype-Tissue Expression (GTEx) project (gtexportal.org). Lung, heart tissue, and whole blood were also included in this analysis (**Extended Data Fig. 4A,B**). We specifically focused on *ACE2*, encoding the first reported receptor for SARS-CoV-2 entry into human cells, neuropilins (*NRP1*, *NRP2*), and the proteases *TMPRSS2*, *FURIN*, Cathepsin B (*CTSB*), and Cathepsin L (*CTSL*), required to cleave the S protein for viral entry and replication^30-33^. Bulk RNA-seq analysis showed a similar expression pattern for *ACE2*, *NRP1*, *NRP2*, *FURIN*, *CTSB*, and *CTSL* in the aorta, coronary, and tibial arteries compared to the lung, with the exception of *TMPRSS2,* which was expressed at lower levels in the arteries (**Extended Data Fig. 4B**).

To investigate the cellular expression of SARS-CoV-2 receptors and entry factors in the human atherosclerotic tissue, we performed an integrated scRNA-seq analysis of human carotid plaques from 10 patients undergoing carotid endarterectomy with atherosclerotic coronary data obtained from 7 coronary samples from 4 heart transplant cases publicly available in GEO (GSE131780; **Fig. 4A**)^34^. The two datasets were merged using the Harmony algorithm, resulting in 16 subclusters of immune cells that corresponded to all major immune populations known to infiltrate human atherosclerotic plaques (**Extended Data Fig. 4C**). The SARS-CoV-2 entry receptors and host entry factors including *NRP1,* a SARS-CoV-2 receptor that can bind FURIN-cleaved S protein to facilitate SARS-CoV-2 viral entry, and the proteases *CTSB* and *CTSL*, were highly expressed in myeloid subclusters, while *ACE2* and the transmembrane serine proteases *TMPRSS2, TMPRSS4, TMPRSS11A*, and *TMPRSS11B* were either undetectable or expressed at low levels (**Extended Data Fig. 4D**). Based on this observation, we subclustered myeloid cells (**Fig. 4B**, **Extended Data Fig. 4E**) and identified two clusters of dendritic cells, three clusters of monocytes/macrophages, one cluster of mixed myeloid cells, and four clusters of macrophages/foam cells that were annotated based on the expression of canonical markers (**Extended Data Fig. 4F**). To identify significant differences in the abundance of myeloid cells between carotid and coronary arteries we performed Milo differential neighborhood abundance testing^35^. This analysis revealed that *TREM2*^high^ macrophages were enriched in coronary tissue, whereas *VCAN*+ monocytes/macrophages and CD1c+ DCs were enriched in carotid samples. *CD16*+ monocytes, inflammatory monocyte/macrophages, *IL1B* DCs, and *CD36*+ mixed myeloid were exclusively present in carotid samples, whereas *LYVE1*+ macrophages were present in coronaries. *SPP1*+ macrophages were present in both tissues (**Fig. 4C**). Overall, *NRP1* was strongly expressed in *TREM2*+, *SPP1*+, *LYVE1*+, and *interferon response genes (ISG)*+*TREM2*+ macrophages, clusters that also expressed *FURIN* (**Fig. 4D**)*. ACE2* and *TMPRSS2* were undetectable in the analyzed myeloid cells.

**Figure 4.**
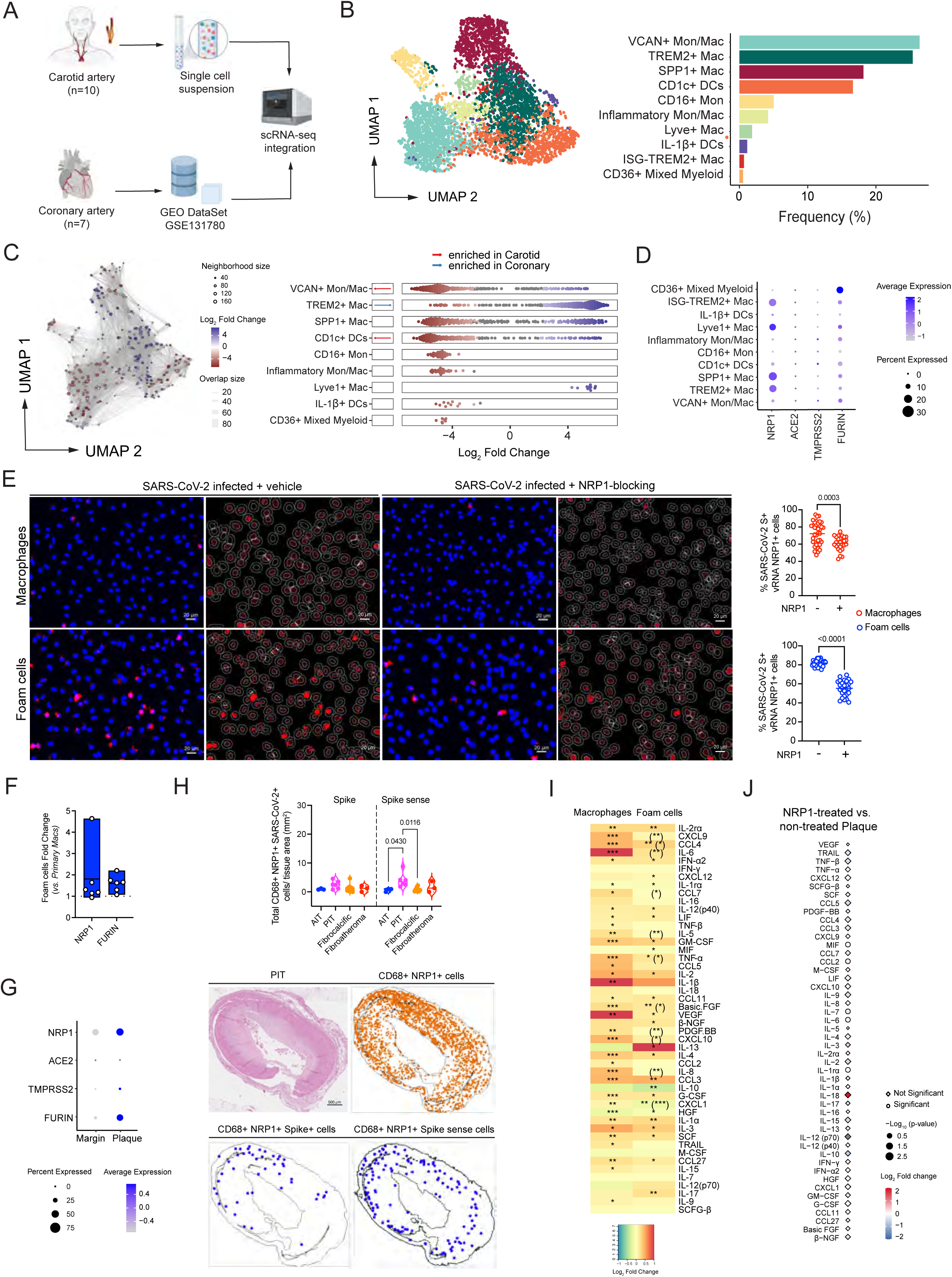
A) Single-cell RNA sequencing of human carotid (*n* = 10) and coronary (*GSE131780*) (*n* = 7) tissue samples. B) UMAP visualization of myeloid cell subclusters from coronary (*n* = 1960 cells) and carotid (*n* = 2900 cells) samples. Bar plot shows the frequency of each myeloid cluster. C) Neighborhood graph of the results from Milo differential abundance testing. Nodes represent neighborhoods, colored by their log2fold change between carotid (*red*) and coronary (*blue*) samples. Non-differential abundance neighborhoods are in white (FDR 10%), and node size reflects total number of cells in each neighborhood. Beeswarm plots show the of log_2_ fold change distribution of neighborhoods between tissue type (FDR 10%). D) Dot plot of the SARS-CoV-2 viral entry factor average gene expression (*color*) and cellular percent of expression (*dot size*) in each myeloid subcluster. E) Representative images of AI-based RNA-FISH quantification showing SARS-CoV-2 spike (S) viral RNA (vRNA) (*red*) and NRP1 (*green*) RNA (*left*), and SARS-CoV-2 antisense (S sense) RNA (*red*) and NRP1 (*green*) RNA (*right*). Quantification of infected macrophages (*top*) and foam cells (*bottom*) with and without NRP1-blocking (EG00229 trifluoroacetate; 100 μM) at 24 hpi. Bar plots show the frequency of NRP1+ SARS-CoV-2 S vRNA+ cells per field *(left)* and NRP-1 SARS-CoV-2 S sense+ cells per field *(right)*. F) Bar plot showing the fold change gene expression of *NRP1* and *FURIN* in uninfected foam cells normalized to primary macrophages (*n* = 6). G) Dot plots of sc-RNAseq results showing the frequency (*dot size*) of cells expressing SARS-CoV-2 viral entry factors colored by average expression in atherosclerotic plaque lesions and paired vasculature margins. H) Violin plot show total CD68+ NRP1+ SARS-CoV-2 vRNA+ normalized by tissue area (mm^2^) in adaptive intimal thickening (AIT), pathological intimal thickening (PIT), fibrocalcific, and fibroatheroma coronaries from COVID-19 patients. Representative images of H&E and spatial analysis of PIT coronary sample showing the location of CD68+ NRP1+ cells (*orange dots*), CD68+ NRP1+ SARS-CoV-2 Spike+ cells (*blue dots, left*) or Spike sense+ (*blue dots, right*) cells. I) Heatmaps of differentially secreted cytokine and chemokine levels from SARS-CoV-2 infected macrophages and foam cells after NRP1-blocking (EG00229 trifluoroacetate; 100 μM). Results are shown as log_2_ fold change between infected and non-treated conditions. Adjusted *P* values < 0.05 were considered significant. Asterisk indicates an adjusted p-value < 0.05 for the comparison of SARS-CoV-2 infected and treated vs infected and untreated. Asterisks in parentheses indicate nominal *P*-value < 0.05 for the comparison between macrophages vs. foam cells. *\*, P < 0.05; **, P < 0.01; ***, P < 0.001*. J) Plot showing the relative expression of secreted cytokines and chemokines from atherosclerotic plaque tissue infected with SARS-CoV-2 and treated with NRP1-blocking *vs.* untreated tissue at 48 hpi. Relative expression is represented in log_2_ fold change colored scale. Statistical significance is expressed as dot size (-log_10_ nominal p-value). Statistically significant values are represented as circles, while not significant changes are represented as diamonds.

As *NRP1* was highly expressed in the *TREM2*^hi^ macrophage populations, identified as foamy plaque macrophages^36^, we used a small molecule that prevents SARS-CoV-2 binding to the B1 domain of NRP-1 and reduces SARS-CoV-2 viral entry^32^, EG 00229, on SARS-CoV-2 infected macrophages and foam cells. NRP-1 blocking reduced the number of SARS-CoV-2 S RNA+ NRP1+ macrophages and foam cells, with the effect ∼ twofold greater in foam cells (**Fig. 4E**). Viral replication was decreased as observed by the reduction in SARS-CoV-2 S sense positive cells (**Fig. 4E; Extended Data Fig. 4G**). *NRP1* and *FURIN* gene expression were higher in uninfected foam cells versus macrophages (**Fig. 4F**) and in the human atherosclerotic plaques from CV patients that did not have COVID-19 compared to paired normal margins (**Fig. 4G**). This indicates that NRP-1 may be preferentially expressed in foam cells and atherosclerotic vasculature, suggesting it would have a key role in mediating SARS-CoV-2 infection of these cells and tissues. Spatial RNA-FISH analysis of human coronary autopsy specimens from COVID-19 patients confirmed that macrophages expressing *NRP1* (CD68+NRP1+ cells) infiltrated coronary lesions, and that these cells expressed SARS-CoV-2 S vRNA and the antisense strand of the S gene, indicating viral replication (**Fig. 4H**). A higher number of NRP1+ macrophages expressing the antisense strand of the S gene were found in PIT coronary lesions, confirming a greater susceptibility of PIT lesions to SARS-CoV-2 infection (**Fig. 4H**).

We next asked whether NRP-1 blocking with EG 00229, which reduces the infection of macrophages and foam cells *in vitro*, would reduce the inflammatory response to SARS-CoV-2. Counterintuitively, EG 00229 triggered a stronger pro-inflammatory response than that elicited by SARS-CoV-2 infection itself, involving the release of several proatherogenic cytokines and chemokines (e.g., IL-6, IL-1β, and CCL2) (**Fig. 4I, Extended Data Fig. 4H**). This response was blunted in foam cells, as shown by the significantly lower release of several pro-inflammatory and proatherogenic cytokines and chemokines versus infected macrophages. Similarly, in SARS-CoV-2 infected human atherosclerotic vascular explants, blocking NRP-1 increased the release of IL-6 and CCL2 (**Fig. 4J**). IL-1β and IL-18 secretion was also increased, but not significantly.

Taken together, these findings suggest that atherosclerotic plaques are particularly susceptible to SARS-CoV-2 infection, and upon infection, a hyperactivated immune response is triggered that could contribute to the increased risk of ischemic cardiovascular events in COVID-19 patients with underlying atherosclerosis. Although blocking the host entry factor, NRP-1, expressed in macrophages and foam cells infiltrating coronary lesions in humans reduced SARS-CoV-2 infection, it also induced a strong inflammatory response, indicating that an alternative strategy is necessary to prevent SARS-CoV-2 infection of coronary vessels and its downstream consequences.

## Discussion

Although SARS-CoV-2 is considered a respiratory virus, COVID-19 patients are at increased risk of cardiovascular complications, including myocardial infarction and stroke. Our study provides the first evidence of SARS-CoV-2 viral presence in human coronary vasculature, and demonstrates viral tropism for vascular lesion macrophages in individuals with severe COVID-19. We found evidence of SARS-CoV-2 replication in all analyzed human autopsy coronaries regardless of their pathological classification, although viral replication was highest in pathological intima thickening (PIT) coronary lesions, early-stage lesions that progress to more advanced atherosclerotic plaques^13,37^. SARS-CoV-2 showed a stronger tropism for the arterial lesions than corresponding perivascular fat, which was related to the degree of macrophage infiltration, consistent with the higher viral replication in PIT lesions and fibroatheromas, atherosclerosis lesions where macrophages are more prevalent. Others have previously reported the presence of SARS-CoV-2 RNA within the heart, the aorta, as well as other distant organs^8,12,38,39^. Our data conclusively demonstrate that SARS-CoV-2 is capable of infecting and replicating in macrophages within the coronary vasculature of patients with COVID-19. Furthermore, SARS-CoV-2 preferentially replicates in foam cells compared to other macrophages, leading to vRNA and viral protein accumulation, suggesting that these cells are more permissive and susceptible to becoming reservoirs of SARS-CoV-2 viral debris in atherosclerotic plaques. Although, viral replication in macrophages and foam cells was abortive, it promoted a strong inflammatory response characterized by release of cytokines implicated in both the pathogenesis of atherosclerosis and increased risk of cardiovascular events such as stroke and myocardial infarction^25,26,40^.

Using an *ex vivo* model of viral infection of human vascular explants, we found that atherosclerotic tissue could be directly infected by SARS-CoV-2, confirming our observation in COVID-19 patient tissue. As in cultured macrophages and foam cells, SARS-CoV-2 infection of vascular tissue triggered an inflammatory response and induced the secretion of key pro-atherogenic cytokines, such as IL-6 and IL-1β. Considering that plaque inflammation promotes disease progression and contributes to plaque rupture, our results provide a molecular basis for how SARS-CoV-2 infection of coronary lesions can contribute to the acute cardiovascular manifestations of COVID-19, including myocardial infarction^3,41^. SARS-CoV-2 infection of coronaries was unrelated to pre-existing clinical characteristics, stage of COVID-19 by illness days, duration of hospitalization at the time of death, or co-morbidities. However, we found a higher accumulation of SARS-CoV-2 S and S antisense vRNA in the coronary vasculature of the three patients with acute ischemic cardiovascular manifestations, including posterior myocardial infarction (P1) and type II myocardial infarction (P2 and P3). Although, evidence of coronary occlusion was not detected at autopsy for two patients with clinical diagnosis of myocardial infarction, these data suggest that SARS-CoV-2 coronary infection may increase cardiovascular risk.

Notably, we observed a similar expression pattern of SARS-CoV-2 receptors and co-factors in human vasculature as that found in the lungs, by systematically analyzing Gtex, a multi-tissue gene expression dataset that includes donors who died from cerebrovascular (> 22%) and heart disease (> 40%). In particular, while *ACE2* expression was low in the aorta and the tibial artery, its expression levels were higher in the coronary artery, comparable to its expression in lung, suggesting that coronary vasculature may be more susceptible to SARS-CoV-2 viral infection than other vascular beds. At the single-cell level, expression of SARS-CoV-2 receptors and factors confirmed our coronary vasculature autopsy findings. Although *ACE2* expression was not detectable by scRNAseq, *NRP1* and *FURIN* were highly expressed in two *TREM2*+ macrophage clusters, known to correspond to plaque foamy macrophages^36^, as well as in clusters of *SSP1*+ macrophages and *LYVE1*+ macrophages. We further found that SARS-CoV-2 infected NRP1+ macrophages in human coronary tissue taken at autopsy, and that viral replication was greater in NRP1+ macrophages present in PIT lesions. Experimentally, a specific inhibitor of the interaction between the b1 domain of NRP-1 and the SARS-CoV-2 S1 CendR^32,33^ reduced SARS-CoV-2 infectivity of human primary macrophages and foam cells, confirming that SARS-CoV-2 infection of macrophage and foam cell is in part neuropilin-1 (NRP-1) dependent. However, an aberrant proinflammatory response associated with NRP-1 inhibition limits the potential therapeutic use of the NRP-1 inhibitor. This proinflammatory effect, although unexpected in the context of SARS-CoV-2 infection, is consistent with previous findings of a protective role of NRP-1 in sepsis and the increased release of pro-inflammatory cytokines (e.g., IL-6) from NRP-1 null macrophages^42^.

Overall, our data demonstrate that SARS-CoV-2 replicates in macrophages within human coronaries of patients who died from severe COVID-19. Our study is limited to the analysis of a small cohort of older individual with COVID-19 and pre-existing atherosclerosis and other medical conditions and comorbidities. Therefore, our observations cannot be extrapolated to younger, healthy individuals. Our study is also limited to cases that occurred during the early phases of the COVID-19 pandemic, and the findings that SARS-CoV-2 replicates in the atherosclerotic coronary vasculature is pertinent only to the viral strains that circulated in New York City between May 2020 and May 2021. Despite these limitations, our study highlights the hyperinflammatory response orchestrated by SARS-CoV-2 infected plaque macrophages and foam cells as a mechanistic link between infection of atherosclerotic coronary vessels and acute cardiovascular complications of COVID-19.

## Acknowledgement

We thank the NYU BSL3 high-containment facility, the NYULH Center for Biospecimen Research and Development, the Histology and Immunohistochemistry Laboratory (CBRD; RRID:SCR_018304), the Experimental Pathology Division (ExPath), the NYU Genome Technology Center (GTC), the NYU Microscopy Laboratory, the NYU Immune Monitoring Laboratory (IML) at the NYU Langone’s Division of Advanced Research Technologies (DART) for their assistance. The DART CBRD, ExPath, IML and the Microscopy Lab cores are supported by the NIH/NCI grant P30CA016087. CBRD is also supported by the Laura and Isaac Perlmutter Cancer Center Support Grant. ExPath is also supported by NIH S10 OD021747. This work was funded by the NIH/NHLBI grant 1R01HL165258 (C.G.). C.G. also acknowledges support from grants NIH/NHLBI R01HL153712, AHA 20SFRN35210252, CZI NFL-2020-218415, U34TR003594. N.E is supported by the AHA Research supplement to promote diversity in science (AHA 965509); M.G.N is supported by the AHA Postdoctoral Fellowship 19-A0-00-1003686; D.D. is supported by the AHA grant 20SFRN35210252; SARS-CoV-2 work in the M.S. laboratory is supported by NIH/NIAID R01AI160706 and NIH/NIDDK R01DK130425; K.A.S is supported by NIH/NIAID 1R01 AI162774-01A1, NYU Grossman School of Medicine start up funds, and NYU Cardiovascular Research Center pilot award. K.J.M. is supported by NIH/NHLBI R35HL R35HL135799 and R01HL084312. The publicly available data used for the analyses described in this manuscript were obtained from the GTEx Portal. The Genotype-Tissue Expression (GTEx) Project was supported by the Common Fund of the Office of the Director of the National Institutes of Health, and by NCI, NHGRI, NHLBI, NIDA, NIMH, and NINDS. The following reagent was deposited by the Centers for Disease Control and Prevention and obtained through BEI Resources, NIAID, NIH: SARS-Related Coronavirus 2, Isolate hCoV-19/USA-WA1/2020, NR-52281; Vero E6-TMPRSS2-T2A-ACE2, NR-54970.

## Author Contributions

Conceptualization, N.E. and C.G.; methodology, N.E., M.G.N., O.S., R.K., K.M, K.A.S., N.N.; BSL-3 experiments N.E., M.G.N, S.J.; other experiments M.S., N.E., O.S, B.C., L.A., D.D., A.V.G.; patient recruitment R.S., S.S., D.F.; clinical data management N.E., S.S., J.N., N.N., R.S., P.F., A.R.; human sample collection, P.F., N.N., A.R., T.M., C.R.; human sample processing N.E., L.A., A.R., D.F., B.C., D.D., data analysis by N.E., D.D., M.G.N., R.K., M.G., D.F., J.N., B.C., D.D.; resources by C.G., K.A.S., A.R.; data visualization by N.E, R.K, M.G.; writing original draft, N.E., C.G.; revision and editing by all authors; supervision by C.G.; project administration and funding acquisition, C.G.

## Extended data- Figure Legends

**Figure S1.**
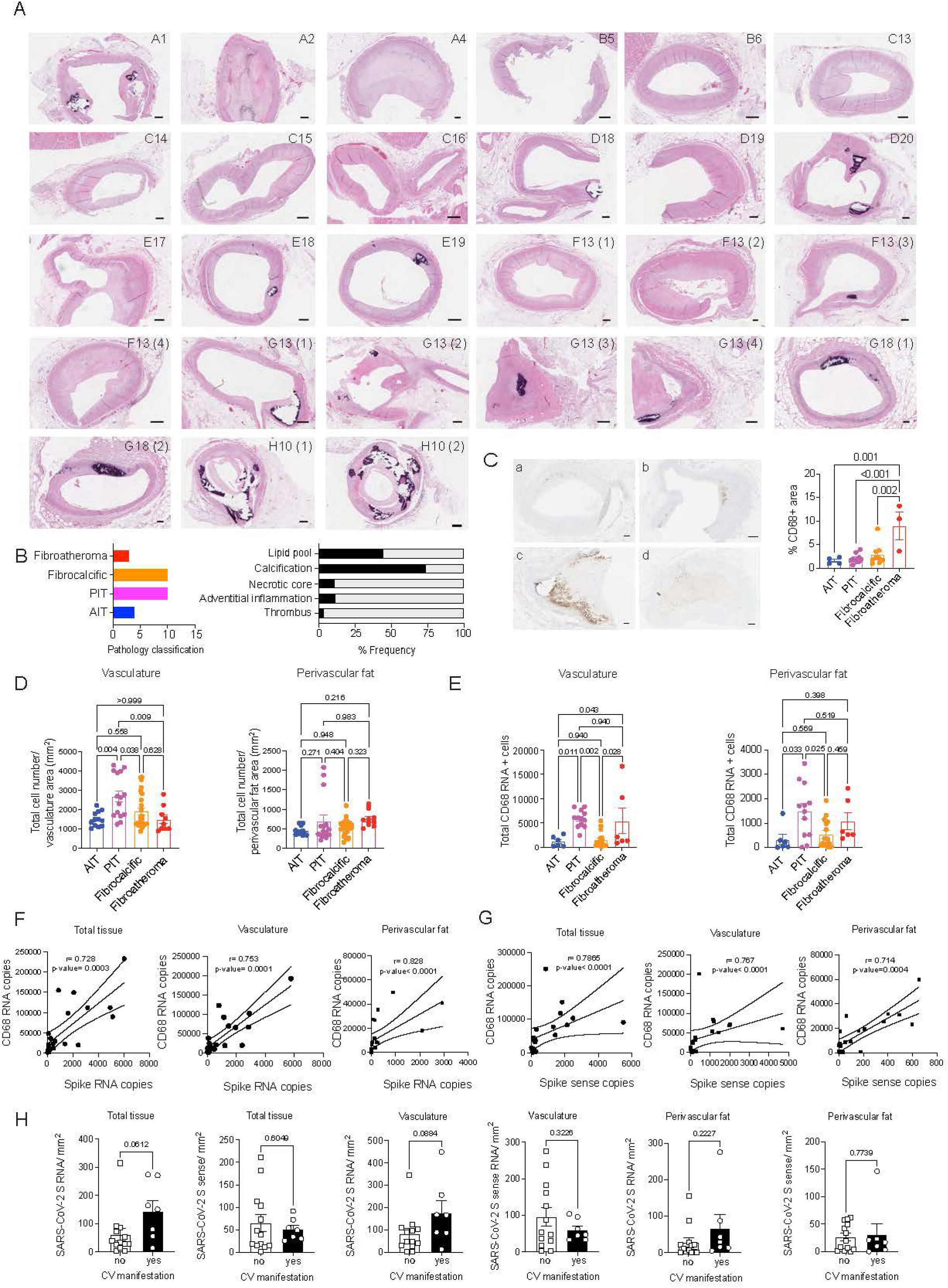
A) Whole scanned hematoxylin and eosin (H&E) images of the 27 coronary samples included in the study. Scale bar = 500 μm. B) Bar plot on the left shows the number of adaptive intimal thickening (AIT; *blue*, *n* = 4), pathological intimal thickening (PIT; *magenta*, *n* = 10), fibrocalcific (*orange*, *n* = 10) and fibroatheroma (*red*, *n* = 3) specimens. The bar plot on the right shows the proportion of coronary tissue sections that presented a positive pathological assessment for lipid pool, calcification, necrotic core, adventitial inflammation, and/ or thrombus. Black bars represent the frequency for each pathological assessment. C) Representative images from CD68 chromogenic immunohistochemistry assays executed in paraffin-embedded coronary specimens classified as (a) AIT, (b) PIT, (c) fibrocalcific, and (d) fibroatheroma of diseased patients with COVID-19 (Scale = 500 μm). Bar plot showing the quantification of % CD68+ area. D) Bar plots of total cell number quantified by AI-based nuclei segmentation normalized by the area (mm^2^) of vasculature and perivascular fat. Each dot represents a tissue section. E) Bar plots of total number of CD68 RNA positive cells quantified in the RNA-FISH in the vascular wall and perivascular fat. Dots represent each quantified tissue section. ANOVA one-way statistical analysis following by post-hoc Tukey’s test for multiple comparisons was performed to evaluate differences between more than two groups. F-G) Scatter plot of Spearman’s rank correlation (95% confidence interval) of total CD68 RNA copies with total SARS-CoV-2 spike (F) and spike sense (G) copies in the whole tissue, vascular tissue, and perivascular tissue specimens. H) SARS-CoV-2 spike (S) and spike sense (S sense) quantification in total tissue, vasculature, and perivascular fat normalized by tissue area (mm^2^) in patients with acute CV complications vs. patients without CV manifestations. Unpaired *t*-test was performed to evaluate differences between two groups.

**Figure S2.**
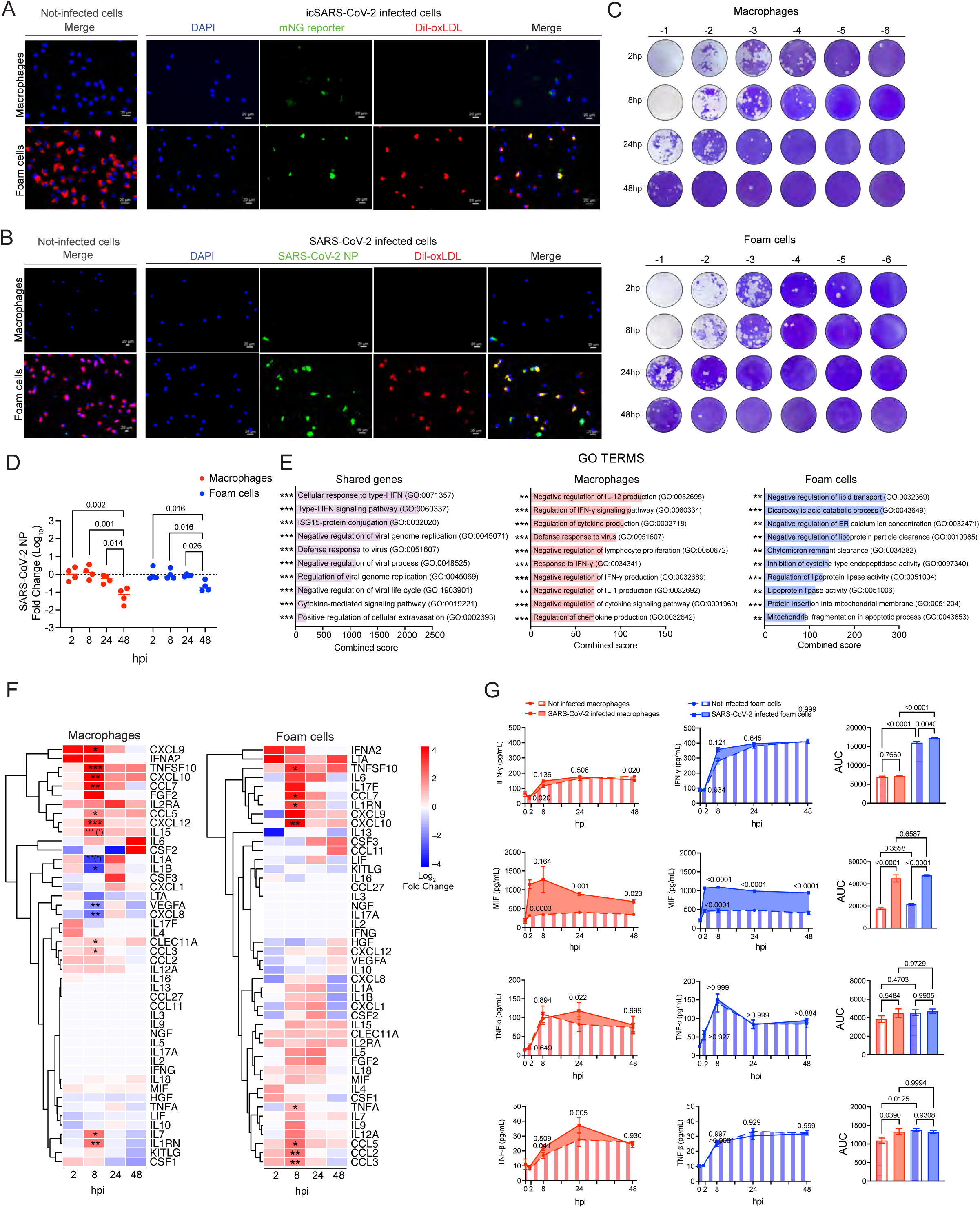
A) Representative fluorescent microscopy images of not-infected (merge) and infected macrophages and foam cells cultured with mNG reporter. B) Representative fluorescence images of not-infected (merge) and SARS-CoV-2 WA1/2020 infected macrophages and foam cells at 24 hours post-infection (hpi). Scale bar = 20 μm. Representative images depict nuclei with DAPI (*blue*), mNG reporter virus or SARS-CoV-2 nucleoprotein (NP) (*green*), Dil-Ox-LDL treatment (*red*), and merged images. C) Representative images of plaque assay in VERO E6-TMPRSS2-T2A-ACE2 cells of culture supernatants of macrophages and foam cells cultured with SARS-CoV-2 USA WA1/2020 at 2, 8, 24, and 48 hpi. Serial dilutions of culture supernatant are represented from left to right (-1 to -6). D) Bar plot showing the Log_10_ Fold change of SARS-CoV-2 viral nucleoprotein RNA levels normalized by 2 hpi samples (*n* = 4) in infected macrophages and foam cells at different timepoints. E) Bar plots showing the combined score of Gene Ontology (GO) enrichment analysis of upregulated genes in infected macrophages (*red*), foam cells (*blue*) and or both macrophages and foam cells (*purple*) vs. non-infected counterparts, ranked by their combined score and selected by significant threshold of *P* < 0.05. *\*, P < 0.05; **, P < 0.01; ***, P < 0.001*. F) Heatmaps of Log_2_ fold change of selected differentially expressed cytokine and chemokine genes in SARS-CoV-2 infected macrophages (*left*) and foam cells (*right*) vs. non-infected counterparts at different hpi. *P*-values were adjusted using Benjamini-Hochberg correction (FDR= 10%). Adjusted *P*-values < 0.05 were considered significant. Asterisk indicates an adjusted p-value < 0.05 for the comparison of infected vs not infected at each timepoint. Asterisk in parentheses (*) indicates an adjusted *P* < 0.05 for the interaction term of the model. *\*, P <0.05*. G) Kinetic plots showing the area under the curve (AUC) of selected cytokines and chemokines in the supernatant of SARS-CoV-2 infected and non-infected macrophages (*red*) and foam cells (*blue*) (*n* = 8). One-way ANOVA statistical analysis followed by Dunnett’s post-hoc for multiple comparisons was performed and *P*-values are denoted for each timepoint. Bar plots represent Mean ± SEM of integrated AUC for each cytokine. One-way ANOVA statistical analysis following Tukey’s multiple comparisons was performed and *P*-values are denoted for group comparisons.

**Figure S3.**
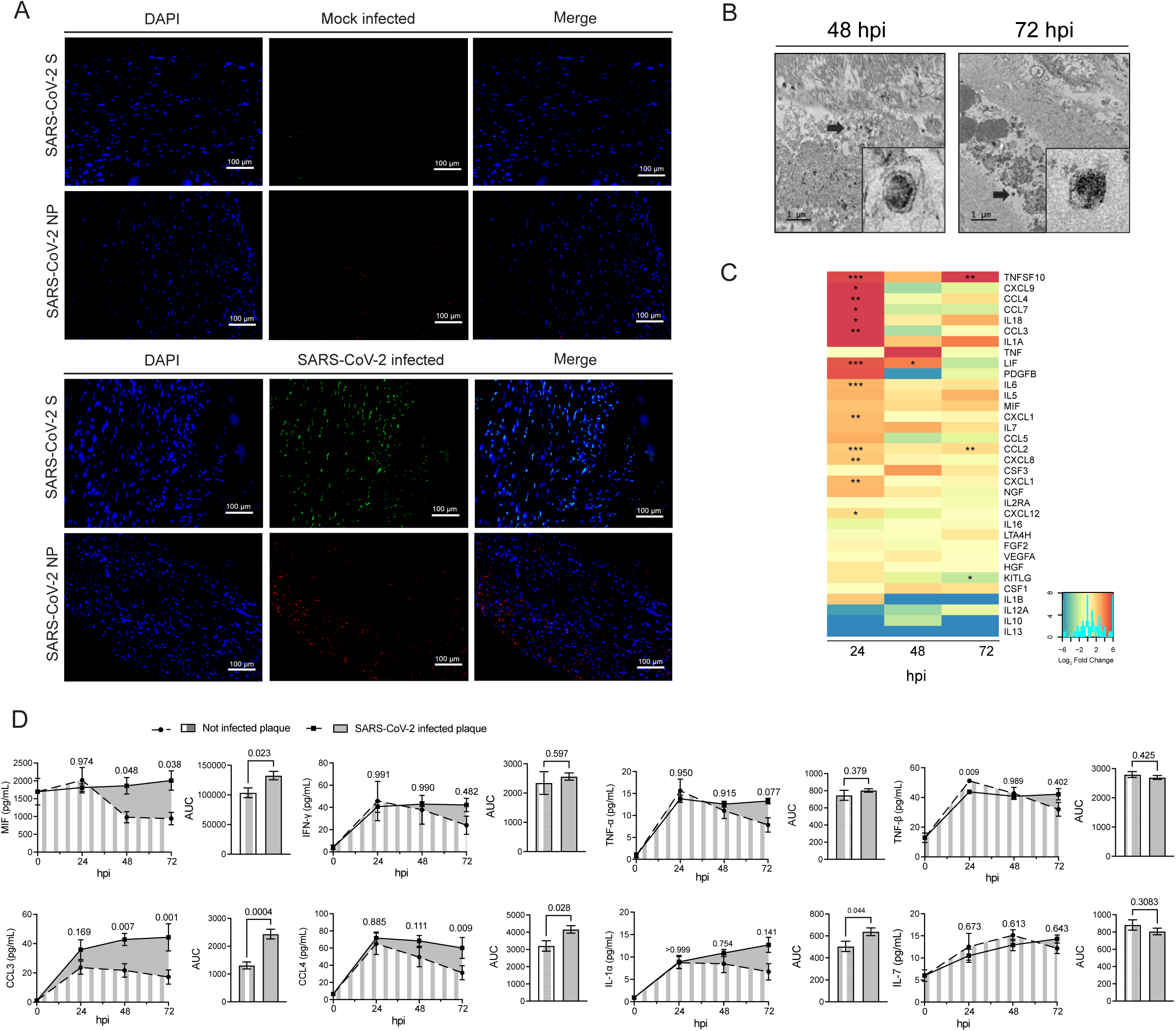
A) Representative immunofluorescence images of human atherosclerotic plaque tissues infected *ex-vivo* with SARS-CoV-2 USA WA1/2020 (10^5^ PFU/mL) (*bottom panel) vs* mock infected control (*top panel*) shows the expression of spike (S) protein and nucleoprotein (NP). Images depict nuclei with DAPI (*blue*), SARS-CoV-2 S protein (*green*) or NP (*red*). Scale bar = 100 μm. B) Electron microscopy of human atherosclerotic carotid plaque tissue infected *ex-vivo* with the SARS-CoV-2 USA/WA1 strain. Scale bar = 1 μm. Black arrows indicate coronavirus-like particles within the carotid atherosclerotic tissue. C) Heatmap of selected cytokine and chemokine genes showing the log_2_ fold changes in SARS-CoV-2 infected carotid vascular explants *vs.* not-infected tissues at different times post-infection. *P* values were adjusted using Benjamini-Hochberg correction (FDR = 10%) and denoted as an asterisk*\*, P < 0.05; **, P < 0.01; ***, P < 0.001.* D) Kinetic plots showing the area under the curve (AUC) of selected cytokines and chemokines secreted by non-infected or SARS-CoV-2 infected carotid vascular explants at different time post-infection. One-way ANOVA statistical analysis following Dunnett’s post-hoc test for multiple comparisons was performed and *P* values are denoted for each timepoint. Bar plots represent Mean ± SEM of integrated AUC for each cytokine. Unpaired *t*-test was performed to compare two groups. *P* < 0.05 was considered significant.

**Figure S4.**
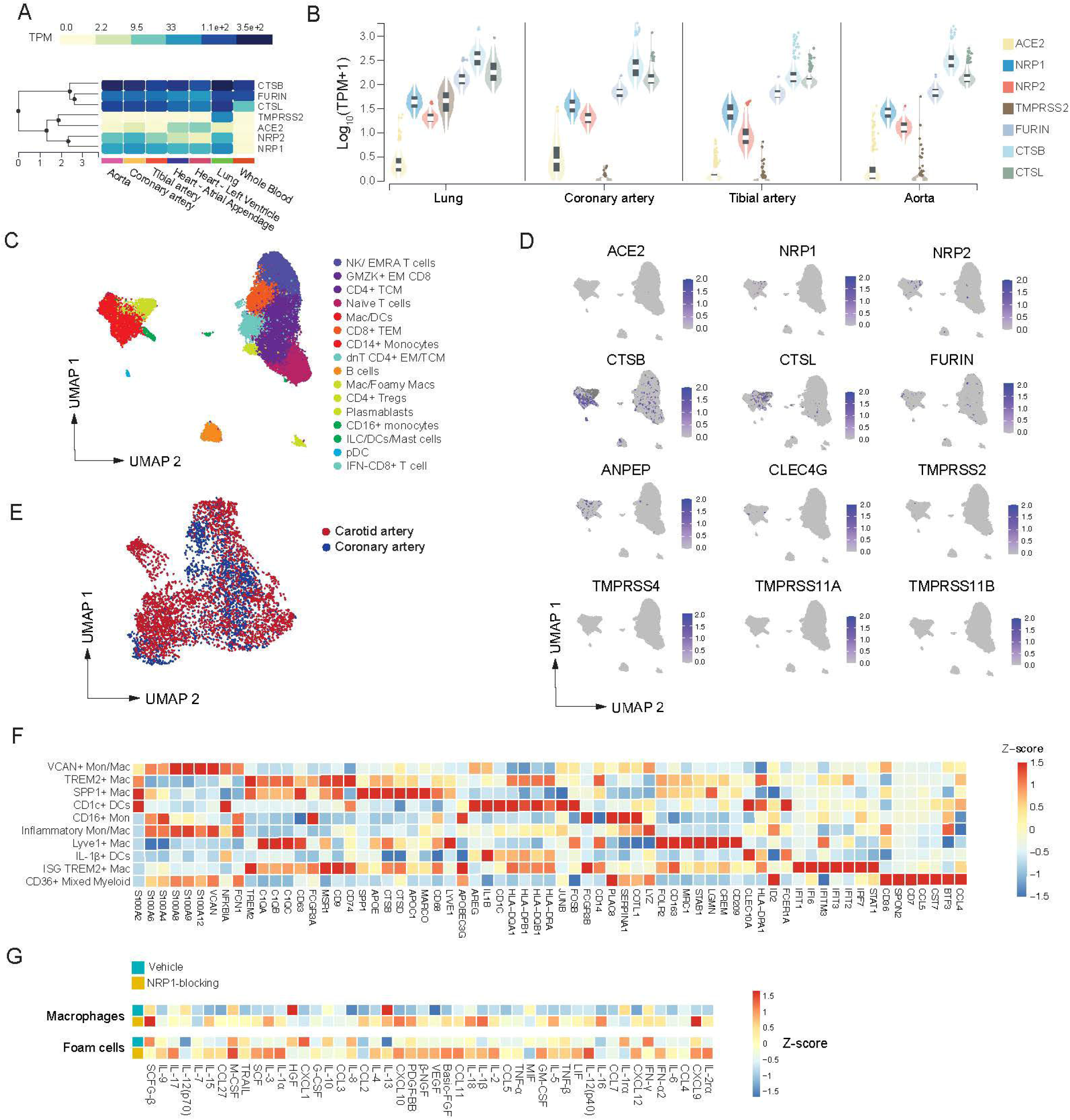
A) Heatmap shows transcripts expression (*TPM*, transcripts per million) of SARS-CoV-2 entry factors (*NRP1, NRP2, CTSB, CTSL, TMPRSS2, ACE2,* and *FURIN*) identified in lung, whole blood, heart (left ventricle and atrial appendage), aorta, and tibial and coronary arteries, using the GTEx public dataset. B) Violin plots showing the Log_10_ TMP of tissue level expression of SARS-CoV-2 entry factors identified using the GTEx public dataset. C) UMAP embedding of integrated total immune cells from carotid and coronary tissue datasets. D) Gene expression of SARS-CoV-2 viral entry factors and related genes projected onto the UMAP of total immune cells. E) UMAP representation of myeloid cell clusters colored by tissue origin. Dots represent individual cells belonging to carotid (*red*) or coronary artery (*blue*). F) Heatmap displaying differentially expressed *z*-score scaled genes (*columns*) across myeloid cell subclusters (*rows*). G) G) Representative images of RNA-FISH showing SARS-CoV-2 spike (S) viral RNA (vRNA) (*red*) and NRP1 (*green*) RNA (left), SARS-CoV-2 S sense (S sense) RNA (*red*) and NRP1 (*green*) RNA (*right*) in infected macrophages (*top*) and foam cells (*bottom*) with and without NRP1-blocking (EG00229 trifluoroacetate; 100 μM) at 24 hpi. H) Heatmap of standardized *z*-scored gene expression of cytokines and chemokines in SARS-CoV-2 infected macrophages and foam cells with or without NRP1-blocking (EG00229 trifluoroacetate; 100 μM).

## Extended data- Supplementary tables

### Tables

Table S1. Demographics and clinical history of COVID-19 patients.

Table S2. Acute clinical manifestations and pathology report information of COVID-19 patients.

Table S3. Histopathology classification and macrophage quantification of post-mortem coronary tissue from patients with COVID-19.

Table S4. Clinical and demographical information of 10 patients undergoing carotid endarterectomy enrolled in the scRNA-seq study.

Table S5. List of antibodies, probes, fluorescent dyes, and reagents.

